# Constraining pesticide resistance using evolution-informed selection regimes

**DOI:** 10.64898/2026.06.02.729543

**Authors:** Luna Qingyang Li, Philip Madgwick, Ricardo Kanitz, Anthony Flemming, Kayla King, Alison Woollard

**Affiliations:** Department of Biochemistry, University of Oxford, Oxford, UK; Syngenta, Jealott’s Hill International Research Centre, Bracknell, UK; Syngenta Crop Protection, Basel, CH; Department of Zoology, University of British Columbia, Vancouver, CA; Department of Microbiology & Immunology, University of British Columbia, Vancouver, CA; Department of Biology, University of Oxford, Oxford, UK

## Abstract

The rapid evolution of pesticide resistance in sexually reproducing pests threatens global food security, yet the evolutionary principles needed to design durable resistance management strategies remain poorly tested experimentally. Theory predicts that deploying multiple pesticide compounds simultaneously should suppress resistance more effectively than sequential rotations, but empirical support in sexual pest populations has remained inconclusive. Here, we directly test how selection regime shapes resistance evolution using a genetically defined, obligately mating *Caenorhabditis elegans* system. We evolved large dioecious populations from a near-isogenic ancestor carrying two major-effect resistance alleles under contrasting pesticide deployment regimes. We show that compound mixtures combined with a substantial refuge consistently produced the strongest constraint on resistance evolution, markedly slowing the spread of resistance even when resistance alleles were neither recessive nor rare. Species-agnostic computational simulations reproduced the overall evolutionary dynamics, suggesting broad applicability. Overall, our results provide direct experimental evidence that pesticide resistance evolution can be predictably constrained by manipulating selection regimes.

## Introduction

The evolution of pesticide resistance in pest populations, particularly insects, threatens global food security and the control of vector-borne diseases. Resistance has been documented in over 600 species^1^, including major agricultural pests and human disease vectors^2^. This results in significant economic damage and loss of human lives^3,4^. Since the introduction of industrial insecticides, resistance has evolved to most chemical classes^5^, and extensive research has characterised its molecular basis^6-17^. However, mechanistic insight alone is insufficient: containing resistance spread in populations exposed to pesticides is fundamentally a problem of population genetics, governed by the underlying evolutionary dynamics. Empirical evidence for managing resistance in dioecious populations is very limited. Insights cannot be directly inferred from work in clonal populations such as in microbial or cancer cells^18,19^ due to the absence of genetic recombination in these systems. Longstanding theoretical work predicts that concurrent application of two compounds as a mixture should outperform rotations^20-24^, assuming no cross-resistance^25,26^, when compounds are efficacious (e.g. in high-dose regimens)^27,28^; some individuals escape exposure in refuges^25,29^; resistance is rare and recessive^29-33^. However, in practice, the optimal strategy for managing pesticide resistance in the field remains a subject of debate, and the use of mixtures continue to be controversial due to limited empirical evidence.

One possible source of empirical evidence comes from aggregated field data. For example, the deployment of crops expressing two *Bt* toxins – effectively a mixture – for the past two decades, suggests that resistance spread could be delayed in some scenarios^34^. Such evidence is heavily influenced by uncontrollable ecological and operational variation in the field, making direct, causal comparisons between strategies difficult. This limitation underscores the need for controlled experimental evolution under laboratory conditions.

Experimental evolution studies of this kind are scarce. A handful of studies have compared resistance management strategies in insect species, including in field trials^35,36^, insect cages^37,38^, and laboratory cultures^39-41^. While the majority report some benefit when more than one compound was used instead of a single compound^36,37,39,40,42-45^, the magnitude of this benefit varied and some studies did not observe this effect^46-48^. Among studies comparing multi-compound strategies, there is likewise no clear agreement on how they should be optimally deployed: some studies support rotations^36,38,48^, whereas others favour mixtures^40,42,45^. Studies involving non-dioecious insects have similarly failed to reach a definitive conclusion^35,49,50^. Overall, the available experimental evidence has not established a relationship between the optimal choice of resistance management strategy and the context in which resistance evolution occurs.

The lack of a clear consensus on managing resistance in dioecious populations can be attributed to several key factors. First, many experiments began with lab-adapted wild ancestral populations previously exposed to pesticides^35,37,38,40,41,45,48,49,51^, introducing uncontrolled genetic variation. In this context, selection acts on heterogeneous backgrounds, making evolutionary trajectories contingent on the founding population. Second, continual, non-plateauing increases in phenotypic resistance suggested a polygenic mechanism^45,51,52^, complicating interpretation and standing in contrast with recent field reports of large-effect monogenic resistance^53-56^. Third, technological constraints of the time prevented accurate tracking of genomic allele frequencies, with resistance inferred indirectly from bioassays or population density^35,37,38,45,48,49,51^. Together, these factors have limited the ability of previous studies to rigorously control the evolutionary context of resistance, precisely characterise its dynamics, and derive broadly generalisable insights.

Here, we establish a novel and experimentally tractable model of dioecious resistance evolution using *fog-2* knockout *Caenorhabditis elegans* to robustly test theoretical predictions on resistance management in animal populations. Although ordinarily androdioecious, *C. elegans* carrying *fog-2* deletion are obligately mating^57,58^, enabling the study of resistance evolution in a fully dioecious population under controlled conditions. Starting from a near-isogenic ancestral population and focusing on two resistance loci of large effect, we robustly test theoretical predictions on the effectiveness of contrasting dual-compound strategies at limiting resistance spread. Through controlled laboratory evolution over 15 generations, we examine whether theoretical predictions concerning the influence of application regimen, compound efficacy, and compound exposure on resistance evolution hold in an experimental system. To assess the generality of these findings, we recapitulate the evolutionary process in a species-agnostic computational simulation, directly comparing simulated predictions with empirical outcomes.

## Results

### Resistance alleles in dioecious *C. elegans* have large effects and no fitness costs

We designed an experimental evolution platform capable of quantitatively assessing the effectiveness of dual-compound strategies across regimen, efficacy, and exposure in a combinatorial design (Fig. 1). Next, we identified suitable resistance alleles in *C. elegans* and generated resistant strains in a dioecious *fog-2* knockout background. Two previously reported mutations, *ben-1* F200Y and *pod-2* A1847V were chosen as they confer large-effect resistance to widely applied pesticides in agriculture – carbendazim and spirotetramat, respectively (Hahnel et al., 2018; Guest et al., 2020). When these resistance mutations were crossed into dioecious *fog-2* knockout strains, they conferred high-level resistance to carbendazim and spirotetramat in a semi-dominant fashion (Figs. 2a, 2b). On carbendazim, the LD_50_ increased from 0.96 µg ml^−1^ in the homozygous susceptible strain to 13.00 µg ml^−1^ in the heterozygote, while no LD_50_ was measurable for the homozygous resistant strain within the limits of solubility (Fig. 2a). A similar pattern was observed for spirotetramat, with LD_50_ values rising from 17.91 to 50.30 µg ml^−1^ in the heterozygotes, and again no measurable LD_50_ for the homozygous resistant strain (Fig. 2b). Additionally, we observed no obvious interaction between these two resistance mutations, with dose-survival data showing no evidence of cross-resistance or synergism/antagonism (Extended Data Fig. 1).

**Fig. 1.**
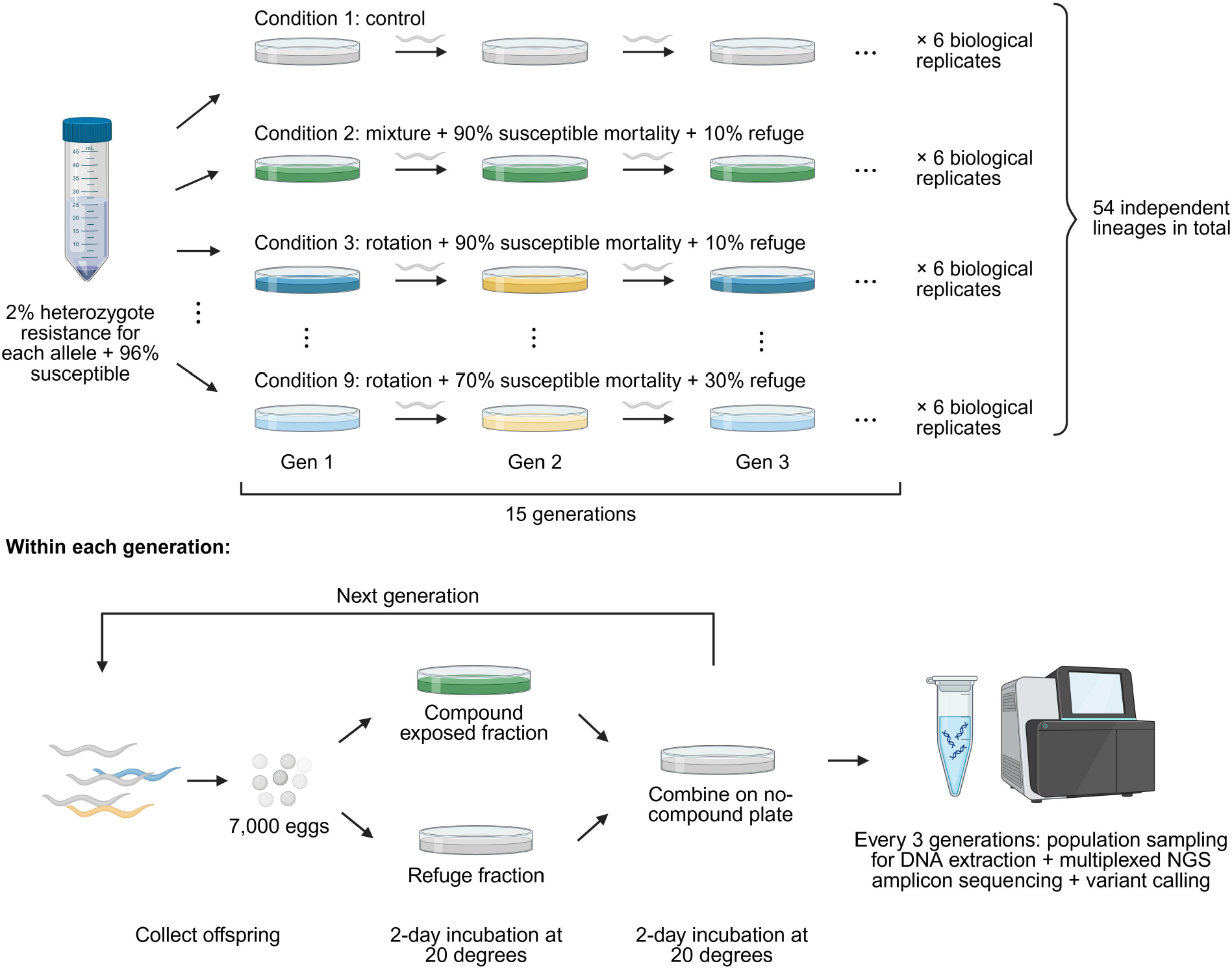
Schematic overview of experimental evolution design and workflow. A total of 54 independent lineages were established, with six biological replicates passaged for 15 generations in each of nine tested conditions. The nine conditions include one no-selection control as well as eight test conditions in a 2 × 2 × 2 factorial design to investigate the effects of regimen choice (mixture vs. rotation), compound efficacy (percentage mortality of susceptible animals), and compound exposure (proportion of population in refuge) on resistance evolution in dioecious populations. 7,000 eggs were seeded to begin each generation, and a sample of the population was taken at the end of generations 0, 3, 6, 9, 12, and 15 to determine population resistance allele frequency using next-generation sequencing.

**Fig. 2.**
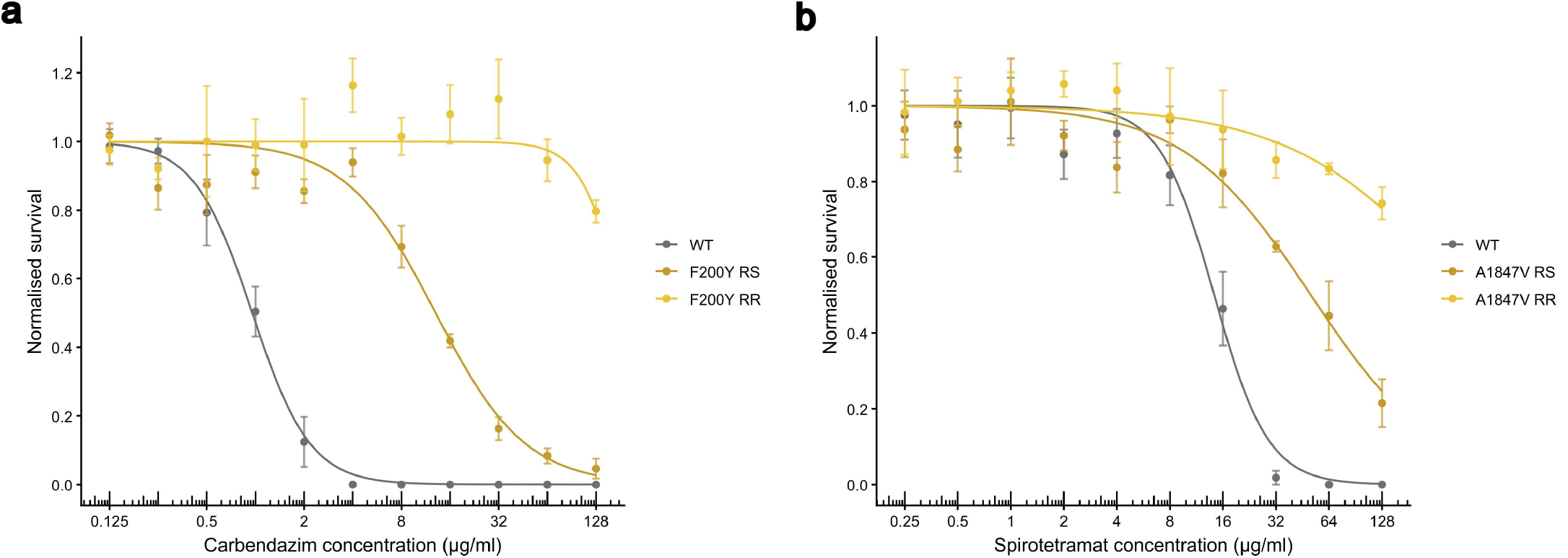
Dose-response analysis of resistance alleles. a) Dose-survival curves showing the effect of harbouring 0, 1, 2 copies of the *ben-1* F200Y allele on resistance to carbendazim. b) Dose-survival curves showing the effect of harbouring 0, 1, 2 copies of the *pod-2* A1847V allele on resistance to spirotetramat. Eggs were seeded onto compound wells on day 0, and survival was quantified as the number of healthy adults after 96h incubation at 20 °C. Data represent control-normalised mean ± SEM from four independent biological replicates (*n* = 4). Curves were fitted using nonlinear regression (two-parameter log-logistic model).

Next, we assessed whether carrying carbendazim or spirotetramat resistance confers a fitness cost. Early-life fitness, relevant for experimental evolution, was assessed using two measures: developmental rate and reproductive capacity. Both measures contribute to early-life fecundity, which is a critical determinant of fitness in experimental evolution studies. Single- and double-homozygous resistant strains were assayed, and none were found to exhibit either defects or benefits in early-life fecundity compared to fully susceptible individuals (Extended Data Fig. 2).

### Selection regime shapes resistance dynamics, favouring mixtures with a larger refuge

Independent lineages originating from a near-isogenic ancestral population carrying ∼2% heterozygous resistance at each locus were evolved for 15 generations under nine distinct selection regimes. The selection regimes, including one no-compound control, were tested for their ability to limit resistance spread. The eight treatment conditions followed a 2 × 2 × 2 factorial design varying regimen choice (mixtures vs. rotations), compound efficacy (90% vs. 70% killing of susceptible individuals), and compound exposure (10% vs. 30% refuge population) (Fig. 1). Compound exposure occurred during the first two days of each four-day generation, allowing mating between exposed and unexposed individuals during the subsequent two days. Chemical concentrations used were calibrated using two-day dose-survival assays (Extended Data Fig. 3). Resistance allele frequencies were quantified in sample populations every three generations by Illumina sequencing.

In all control replicates, both resistance alleles fluctuated around their initial frequencies (Extended Data Fig. 4), confirming that they are neutral under laboratory conditions. Evolution on compound produced distinct outcomes depending on the choice of selection regime (Fig. 3). Increasing the size of the refuge from 10% to 30% of the population was found to be highly effective at limiting resistance spread in all conditions tested, especially for mixtures. Overall, both resistance alleles exhibited similar evolutionary dynamics when subjected to the same selection regime.

**Fig. 3.**
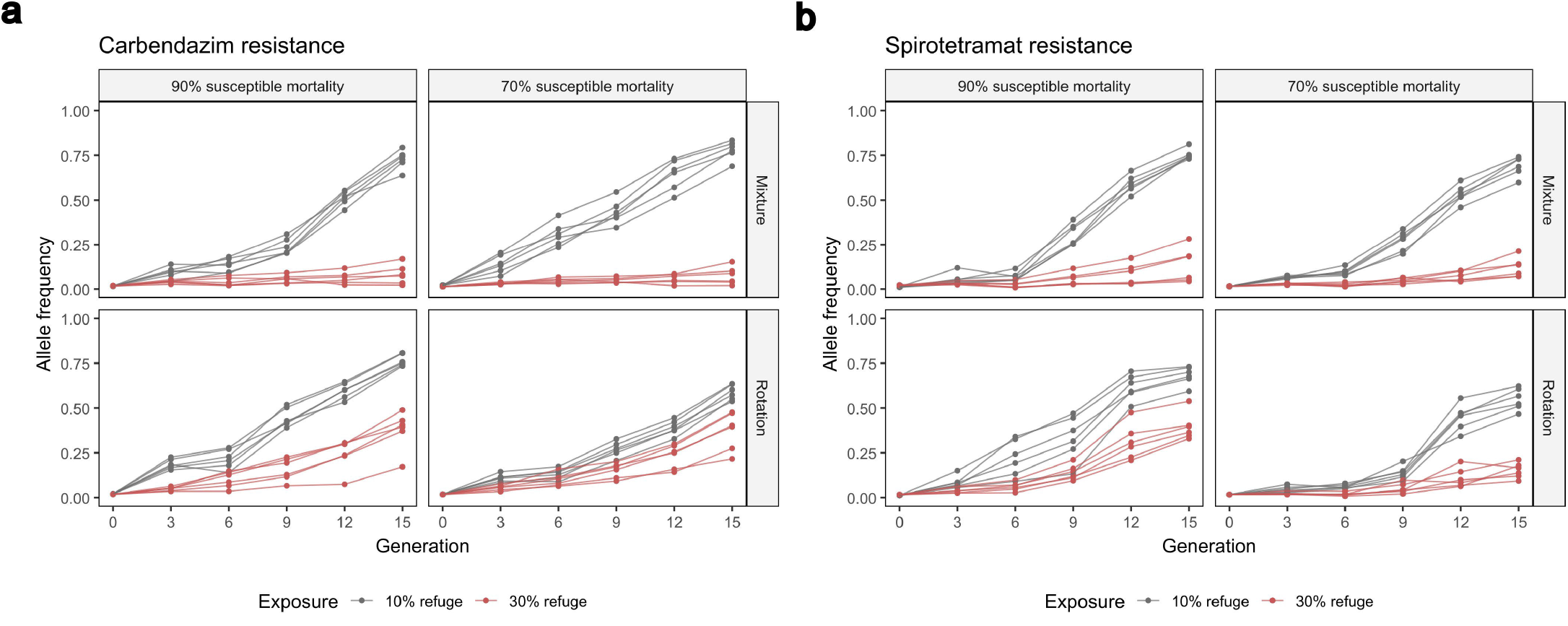
Resistance allele frequency dynamics under different selection regimes. a) Change in frequency of the carbendazim resistance allele. b) Change in frequency of the spirotetramat resistance allele. Allele frequencies were measured at the end of every three generations using high-depth Illumina amplicon sequencing. Data represent individual biological replicate trajectories across 15 generations of experimental evolution (*n* = 6).

To quantify differences in evolutionary outcomes, we derived selection coefficients for individual biological replicates by logistic-growth curve fitting (see Methods), and compared their values across selection regimes (Figs. 4a, 4b). Mixtures with a larger refuge (30%) had selection coefficients three-to four-fold lower than the least effective strategies, indicative of vastly reduced resistance spread. Surprisingly, such favourable outcomes were largely insensitive to compound dose, with similar rates of resistance spread across both compound efficacies tested. This suggests that reduced-dose regimens have the potential to effectively limit the spread of existing resistance. When comparing reduced-dose mixtures with full-dose rotations of similar selective strengths (91% and 90%, respectively), mixtures performed at least as well for the same refuge size, with a larger refuge further favouring mixtures.

**Fig. 4.**
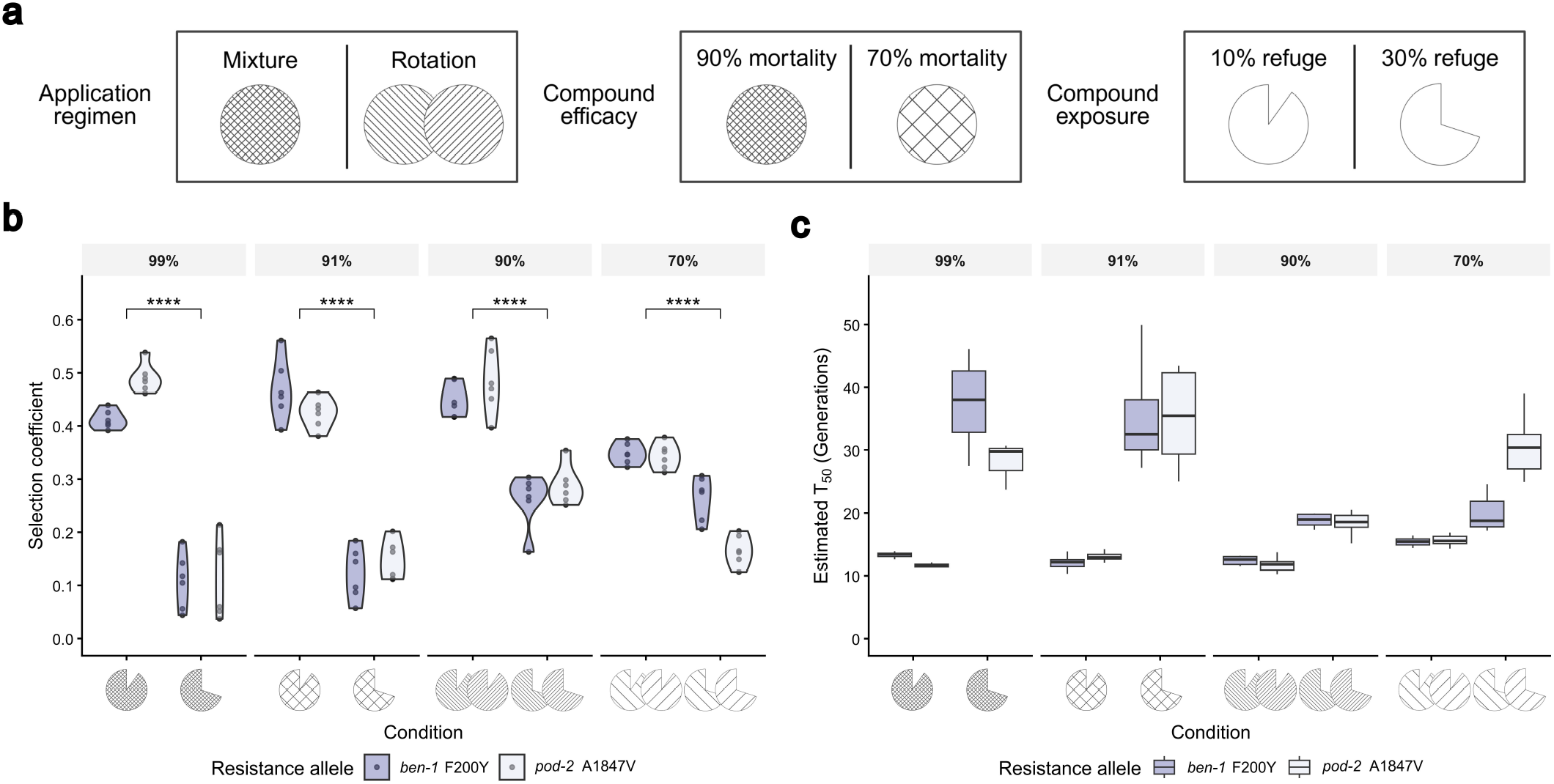
Resistance allele selection coefficient and estimated T_50_. a) Schematic representation of symbols used in panels b and c. b) Selection coefficient for each replicate (shown as dots) derived from experimental evolution data using a logistic growth model. Selection regimes are grouped by total compound efficacy (percentage mortality of susceptible animals not in refuge), as indicated in the top panel (99%, 91%, 90%, 70%). Within all efficacy groups, increasing the refuge proportion from 10% to 30% significantly reduced the selection coefficient (Two-way ANOVA, p = 1.76 × 10^-13^, p = 1.96 × 10^-13^, p = 2.76 × 10^-9^, p = 8.73 × 10^-10^, from left to right). *n* = 6. c) Estimated time to 50% allele fixation (T_50_) based on derived selection coefficients, assuming each resistance allele begins in the population at 1% abundance. Data represent outlier-excluded boxplots across biological replicates (*n* = 6).

To translate selection coefficients into a practically relevant metric, we used experimentally derived estimates to computationally predict the time required to reach 50% resistance (T_50_) (see Methods), assuming an initial allele frequency of 1% (Fig. 4c). Mixtures with a larger refuge, the optimal strategy, doubled and in some cases quadrupled T_50_ relative to alternative regimens. However, due to the non-linear relationship between selection coefficient and T_50_, small differences in selection coefficient generated large variation in T_50_. Therefore, these estimates should be interpreted cautiously.

### Simulations recapitulate experimental outcomes but overestimate selection strength

To assess the generalisability of the experimental results, we developed a species-agnostic simulation of the underlying evolutionary dynamics. The model adopts Wright-Fisher dynamics with discrete generations and is parameterised using baseline resistance and fitness measurements from single-generation assays. The same eight resistance management strategies were simulated as in the laboratory experiments.

Overall, simulation outcomes resembled experimental results (Extended Data Fig. 5), supporting the generalisability of our findings to dioecious pest populations. Increasing refuge size remained effective at limiting resistance spread, particularly in mixtures, mirroring what was observed experimentally. Across comparable strategies, lowering efficacy did not increase resistance spread in most cases, consistent with the experimental observation that reduced-dose regimens can achieve similar control with lower environmental impact. The exception was mixtures with a larger refuge, where simulations showed increased resistance spread at lower compound efficacy, in contrast to laboratory evolution. The sensitivity of this selection regime to dose therefore warrants further investigation.

To quantify discrepancies between experimental and simulated outcomes, selection coefficients were fitted to the mean trajectory of each simulated condition, as for the experimental data. Simulations generally overestimated selection coefficients, particularly for carbendazim resistance (Fig. 5a) relative to spirotetramat resistance (Fig. 5b), suggesting a compound-specific explanation. Overestimation was most pronounced when the selection pressure was the strongest. These discrepancies show that multigenerational evolutionary dynamics cannot be fully inferred from single-generation data alone, as in the simulation.

**Fig. 5.**
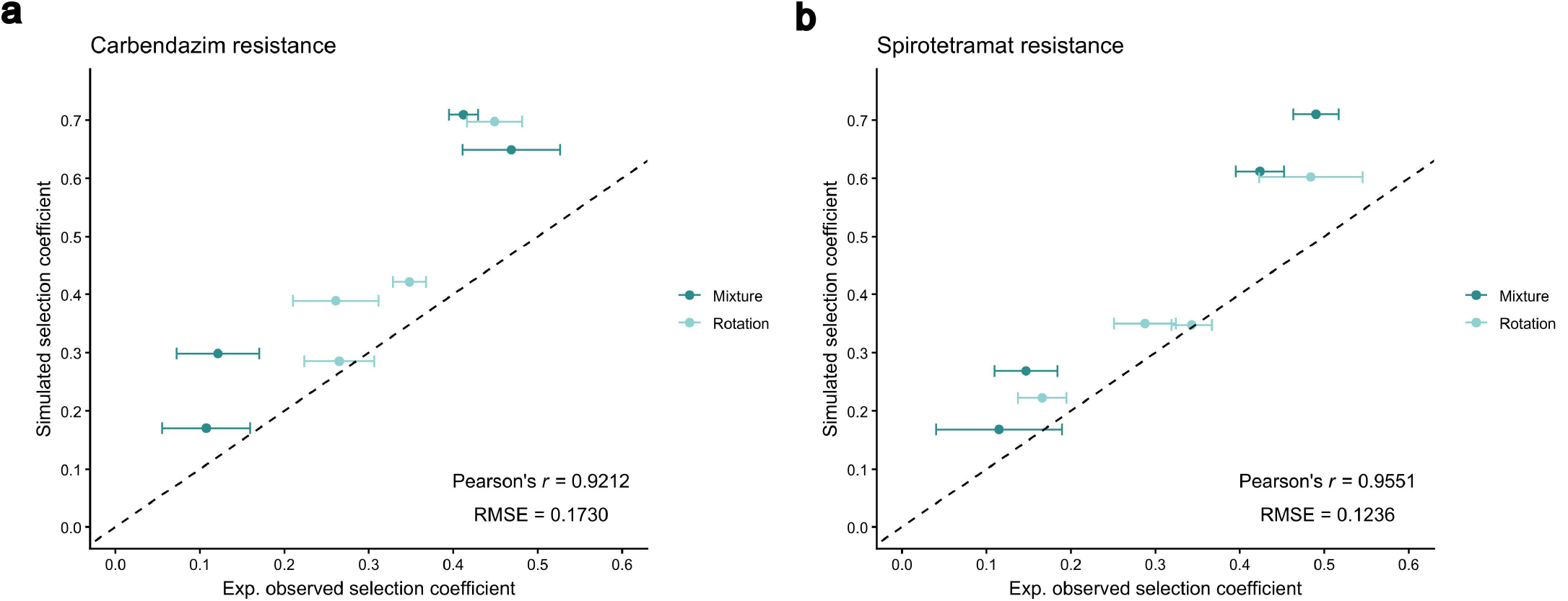
Comparison of experimentally derived and computationally predicted selection coefficients. a) Carbendazim resistance. b) Spirotetramat resistance. Data represent the mean ± SD of experimentally derived selection coefficients (*n* = 6) on the x axis, plotted against the fitted selection coefficient of the mean trajectory from 500 stochastic computational simulations on the y axis. Pearson’s *r* and root mean square error (RMSE) were calculated using the experimental mean. The dotted line indicates equivalence where x=y.

## Discussion

Despite extensive theoretical work, the optimal resistance management strategy in dioecious pest populations lacks robust empirical support. We constructed an experimental evolution system in obligately mating *C. elegans* to directly compare dual-compound resistance management strategies. We found that mixtures with a larger refuge most effectively suppressed the spread of resistance from the same ancestral population. This finding corroborates longstanding theoretical predictions emphasising the importance of refuges for the effective deployment of mixtures^20-24,33,34,61^. By maintaining a fraction of the population free from selection, refuges preserve susceptible alleles which can be reintroduced during sexual reproduction, slowing the formation of double- or homozygous-resistant genotypes. Mixtures can further suppress resistance because partially resistant individuals may still be eliminated through incomplete dominance, or through the alternative compound via the “redundant kill” effect 23, whereas rotations cannot exploit this mechanism. Moreover, while theory suggests that mixtures are the most effective when resistance is recessive and initially rare^22,24,29-32,34^, our findings indicate that refuge size exerts a far stronger influence than either of these factors. Notably, mixtures with a larger refuge outperformed rotations even when resistance alleles were neither recessive nor rare. This has important practical implications, as refuge size can be readily manipulated in field settings. Mixtures may nevertheless fail when cross-resistance occurs^25,26,36^, which has so far been difficult to quantify in natural populations^62^ or characterise in the laboratory^63-65^. The experimental framework developed in this study provides a tractable system for investigating how such cross-resistance traits arise and spread in dioecious populations. Indeed, the rapid generation time of *C. elegans*, combined with the capacity to breed large populations^66^, stablishes this organism as an exceptionally powerful system for investigating pesticide resistance evolution.

Our computational simulations broadly predicted experimental evolutionary outcomes, but their accuracy varied between compounds and with selection strength. Predictions were more accurate for spirotetramat than for carbendazim, likely reflecting differences in their modes of action. Carbendazim, a β-tubulin inhibitor, has been shown to induce paralysis in *C. elegans* from L2 onwards at sublethal doses^67^, whereas spirotetramat primarily affects early developmental stages with limited impact on adults^59,68^. Consequently, carbendazim sensitivity may impose fitness costs that extend beyond survival to adulthood, potentially affecting behaviour and reproductive success in ways not captured by predictions relying solely on survival parameters. Behavioural components of fitness and resistance^48,69-71^, though more difficult to quantify, may therefore be important for accurately predicting resistance trajectories. Discrepancies between simulation and experiment were most pronounced under strong selection, where small deviations in allele frequency early in the process can strongly influence long-term evolutionary outcomes^72^. These results show that computational models complement, but cannot replace, experimental studies of resistance evolution. Although simulations can rapidly explore a broad range of scenarios at scales that are experimentally infeasible^73-81^, their predictive power depends on how accurately they capture the underlying biology. Thus, although simulations capture broad evolutionary trends, they are less able to predict more nuanced effects, highlighting the need for multigeneration laboratory evolution studies.

Our results broadly suggest that the logic of constraining resistance evolution in dioecious populations differs fundamentally from that in clonal systems. In antivirals and cancer therapeutics where clonal populations of resistance dominate, compound combinations are widely used^82-85^. Since resistance in clonal systems typically must arise from *de novo* mutations, combinations act to reduce population size and thus mutation supply, thereby substantially delaying the emergence of fully resistant genotypes^86-90^. Resistance evolution in dioecious populations such as pest insects is less constrained by the initial emergence of resistance alleles. This is because recombination maintains high standing genetic variation^91-93^, and resistance alleles may already exist prior to widespread pesticide use^94^. Crucially, we show that resistance control in dioecious populations can be achieved after its emergence, during the selection phase. Unlike in clonal systems where a fully resistant genotype is expected to rapidly sweep through the population under selection^95-98^, we demonstrate that it is possible to constrain the spread of resistance in dioecious populations by altering the selection regime. Mixtures with larger refuges remain effective even when resistance to both components is present – with the selection regime outweighing the molecular mechanism of resistance as the major determinant of resistance spread.

In conclusion, our results demonstrate that resistance evolution in dioecious animals is strongly shaped by selection regime. Although this study centres on the selection-phase dynamics of monogenic resistance, it establishes a foundation for examining how selection interacts with the emergence and inheritance of more complex resistance traits. By establishing clear, predictable links between selection structure and evolutionary outcome, this work provides robust empirical evidence that resistance management in dioecious systems can be guided by evolutionary principles, with implications for pest and disease vector control. These findings highlight the potential for evolution-informed resistance management strategies to support sustainable agricultural practices, ultimately balancing long-term control with wider environmental and economic considerations.

## Methods

### Strain construction in dioecious *Caenorhabditis elegans*

Genetic crosses were performed between the dioecious strain BS553 *fog-2*(oz40) and homozygous carbendazim- or spirotetramat-resistant lines to generate resistant mutants in a dioecious background. Specifically, BS553 was separately crossed with the *ben-1*(F200Y) strain (Hahnel et al., 2018) and the *pod-2*(A1847V) strain (Guest et al., 2020). Males obtained from the resistant strains by heat shocking L4 larvae were subsequently mated with BS553 females isolated prior to adulthood.

F_1_ offspring, which contain hermaphrodites, were allowed to self-fertilise on compound-treated plates (10 µg ml^-1^ carbendazim or 40 µg ml^-1^ spirotetramat). F_2_ individuals that reached the L4 stage under compound exposure, consistent with at least heterozygous resistance, were isolated into single wells of a 12-well plate. Females unable to produce eggs without mating, therefore homozygous for *fog-2*(oz40), were selected and backcrossed with non-resistant BS553 males. Progeny, homozygous for *fog-2*(oz40), were propagated on compound-treated plates under the same conditions for 3 – 4 generations. A single male-female pair from the final generation was isolated and propagated on a separate plate. The resistance genotypes of the parental worms were confirmed by Sanger sequencing, to establish stocks of obligately mating strains resistant to carbendazim (AW2501) and spirotetramat (AW2502).

### Dose-response assays

Dose-response assays were performed by pipetting approximately 100 worm eggs, obtained by population bleaching (Sulston & Hodgkin, 1988), to a twofold dilution series of carbendazim (0.125 – 128 µg ml^-1^) or spirotetramat (0.25 – 128 µg ml^-1^) on NGM plates seeded with *E. coli* OP50. The solvent used for all treatments was a 1:1 mixture of water and isopropanol, with compounds added to plates before melted NGM agar. Plates were incubated at 20°C for 96h, and survival was scored as the number of healthy worms that developed to adulthood. Each assay included at least three biological replicates. Dose-survival curves were fitted using control-normalised survival data and a two-parameter log-logistic model implemented with the *nls* package in R.

For 2-day dose-response assays, chemical compounds were added to wells at their final concentrations (4 – 9 µg ml^-1^ for carbendazim; 24 – 50 µg ml^-1^ for spirotetramat). After 48h of incubation at 20°C, worms were transferred to no-compound plates by chunking the entire well, and survival was scored at 96h by counting the number of healthy worms that developed to adulthood. Curve fitting was performed as described above.

### Baseline fitness assays

Developmental timing was measured by pipetting approximately 75 worm eggs, obtained by population bleaching, onto 12-well plates incubated at 20°C in the absence of compound. The time (in hours) at which the first egg was laid in each well was recorded.

Reproductive capacity was assayed by pipetting approximately 75 worms eggs, obtained by population bleaching, onto 12-well plates incubated at 20°C in the absence of compound. Plates were incubated for 96h, after which one randomly chosen mature female from each well was spot-bleached by adding 20µl of 5% sodium hypochlorite solution, and the number of intact eggs carried by each individual was counted. Twelve biological replicates were performed for both developmental timing and reproductive capacity assays.

### Experimental evolution

At the beginning of generation 0, 60 homozygous resistant eggs (30 carrying each resistance allele from the strains AW2501 and AW2502) and approximately 2,940 susceptible BS553 eggs were seeded together on each plate by pipetting the appropriate volume following population bleaching. This yielded an initial frequency of 1% for each resistance allele on 54 starting plates. No selection was applied during generation 0. At the end of generation 0, six plates were randomly pooled to form the starting population for each experimental condition, thus averaging out allele frequency variation between plates. These nine founding populations were considered nearly isogenic apart from the resistance loci. A 10% sample (of the washed off adult population) from each of the nine generation 0 pools was collected and frozen for allele frequency analysis. These samples were gently centrifuged to remove bacteria in the supernatant and stored in 150µl TE buffer at -80 °C to protect DNA integrity.

The remainder of each pooled population was then bleached, and six biologically independent lineages of approximately 7,000 eggs each were established to begin selection in generation 1. Nine experimental conditions were tested in total: one control (no compound selection) and eight treatment conditions differing in (i) regimen (mixture vs rotation), (ii) individual compound efficacy (90% vs. 70% mortality), and (iii) total compound exposure (10% vs. 30% refuge). Regimen was implemented by applying the appropriate compound(s) in each generation: mixture treatments received both carbendazim and spirotetramat, whereas rotation treatments alternated between carbendazim in odd generations and spirotetramat in even generations. Compound efficacies were derived from 2-day dose-response data for the susceptible BS553 strain: 7.7 µg ml^-1^ (90% mortality) and 6.1 µg ml^-1^ (70% mortality) for carbendazim; 46 µg ml^-1^ (90% mortality) and 33.5 µg ml^-1^ (70% mortality) for spirotetramat. Refuge size was controlled by changing the number of eggs seeded on no-compound vs. compound-treated plates immediately after bleaching. A 10% refuge consisted of 700 eggs pipetted onto 5.5 or 9cm plates (so as not to exhaust the food supply) with no compound, whereas this was increased to 2,100 eggs for 30% refuge conditions. For the compound exposed fractions, the remainder of the 7,000 eggs to be taken forward to the next generation were pipetted on to 3cm plates containing the appropriate compound concentrations.

All plates were incubated at 20°C for 48h. After incubation, worms from compound-treated plates were transferred to their corresponding no-compound (refuge) plates by chunking the entire 3cm plate. Once worms had migrated off the agar chunks (∼6h), the chunks were removed, and populations were incubated for an additional 48h at 20°C to allow mating. Four days after the generation began, worms were washed off plates into individual tubes. At every three generations (generations 3, 6, 9, 12, 15), approximately 10% of each population was sampled and frozen for allele frequency analysis. Collected samples were gently centrifuged to remove bacteria in the supernatant and stored in 150µl TE buffer at -80°C to protect DNA integrity. Populations were bleached at the end of every generation to obtain 7,000 eggs for the next generation. This procedure was repeated for 15 generations.

### Resistance allele frequency analysis from sampled populations

To determine resistance allele frequencies in evolving populations, we extracted pooled genomic DNA from all frozen samples taken at three-generation intervals between generation 0 and 15. Worm population samples in TE buffer were thawed from −80□°C storage, and DNA was extracted using the Qiagen DNeasy Blood & Tissue Kit, according to the manufacturer’s instructions.

Resistance loci were amplified by barcoded PCR for multiplexed amplicon sequencing. PCR amplification of target loci with combinatorial custom barcodes (barcode sequences in Supplementary Table□1) was performed using high-fidelity Q5 polymerase (NEB) in 25□µL reactions containing 0.8□µL of each primer (20µM). For each locus, six unique forward primers were used to distinguish samples from each biological replicate, and five unique reverse primers were used to distinguish samples from different generations (3, 6, 9, 12, 15). Altogether, for each of the nine conditions, this produced a library of 30 multiplexed amplicons per locus, 60 amplicons per library. The generation 0 samples (18 amplicons in total) were amplified in a standalone library to increase the per sample read depth, capturing low resistance allele frequencies in the starting populations. For the *ben-1* locus, PCR cycling conditions were 98°C 30s, 32 cycles of 98°C 10s, 62.2°C 30s, 72°C 30s, followed by a final extension at 72°C for 2min. For the *pod-2* locus, cycling conditions were 98°C 30s, 30 cycles of 98°C 10s, 62.4°C 30s, 72°C 30s, followed by a final extension at 72°C for 2min.

The integrity of each amplicon (∼240bp) was confirmed by visualising 5µl of the PCR product on 1% agarose gels. Within each library, the remainder of the PCR products were pooled and purified using the NEB Monarch PCR & DNA Cleanup Kit, according to the manufacturer’s instructions. DNA concentration in each pooled, cleaned-up library was measured using a Nanodrop (DeNovix DS-11 FX+), and all ten libraries were sequenced using paired-end reads 150bp in length (PE150) on an Illumina NovaSeq at a commercial facility (Eurofins Genomics).

A custom bioinformatic pipeline was constructed to analyse allele frequencies from the NGS data. Briefly, sequencing reads passing quality control were demultiplexed by forward and reverse barcodes, aligned to the relevant reference sequence, and variants were called. On average, each demultiplexed sample contained at least 10,000 reads over each resistance locus, allowing the identification of allele frequencies as low as 1%. Resistance allele frequency was calculated as the fraction of reads supporting the resistance allele. The command-line pipeline used for bioinformatic analysis is available at https://github.com/LunaL49/DioeciousEvol/tree/main/bioinformatics.

### Estimation of selection coefficient and time to 50% resistance

The selection coefficient (*s*) for each biological replicate was estimated by fitting a logistic growth model of the resistance allele frequency^100^:

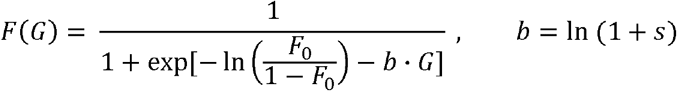

where *F(G)* is the resistance allele frequency at generation *G, F* _0_is the initial resistance allele frequency at generation 0, and *b* is the fitted parameter related to the selection coefficient. Fitting was performed with *b* as the only free parameter using the *nlsList* function from the R package *nlme*.

Time to 50% resistance (T_50_) was predicted from the fitted selection coefficient assuming an initial resistance allele frequency (F_0_) of 1% using the continuous-time approximation (Madgwick & Kanitz, 2022a):

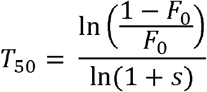

#### Computational simulation

A discrete-generation Wright-Fisher model was implemented to simulate the dynamics of resistance alleles under experimental selection. Within each generation, the simulation proceeds as follows:

1. Generate a population with the current allele frequency distribution.
2. Apply selection to each genotype.
3. Perform random dioecious mating to produce the genotypes of the offspring generation.
4. Incorporate stochastic multinomial sampling to simulate genetic drift.
5. Record genotype frequencies in the offspring generation.

Both resistance alleles were initialised at 1% frequency, exclusively in heterozygotes. Model parameters, including resistance recovery and dominance coefficients, were derived from empirical 2-day dose-response assays. Population size was held constant across generations. Simulations were run using R, with code available at https://github.com/LunaL49/DioeciousEvol/tree/main/simulation. Selection coefficients were estimated in the same way as from experimental data.

## Supporting information

All supplementary figures and legends

## References

1 Mota-Sanchez, D. & Wise, J. C. The Arthropod Resistance Database, <http://www.pesticideresistance.org> (2025).

2 Ranson, H. & Lissenden, N. Insecticide Resistance in African Anopheles Mosquitoes: A Worsening Situation that Needs Urgent Action to Maintain Malaria Control. Trends Parasitol 32, 187–196 (2016). 10.1016/j.pt.2015.11.010

3 Bradshaw, C. J. et al. Massive yet grossly underestimated global costs of invasive insects. Nat Commun 7, 12986 (2016). 10.1038/ncomms12986

4 World Health Organization. World malaria report 2025. (World Health Organization, Geneva, 2025).

5 Sparks, T. C. & Nauen, R. IRAC: Mode of action classification and insecticide resistance management. Pestic Biochem Physiol 121, 122–128 (2015). 10.1016/j.pestbp.2014.11.014

6 Siddiqui, J. A. et al. Insights into insecticide-resistance mechanisms in invasive species: Challenges and control strategies. Front Physiol 13, 1112278 (2022). 10.3389/fphys.2022.1112278

7 Bass, C., Denholm, I., Williamson, M. S. & Nauen, R. The global status of insect resistance to neonicotinoid insecticides. Pestic Biochem Physiol 121, 78–87 (2015). 10.1016/j.pestbp.2015.04.004

8 Guo, L. et al. Convergent resistance to GABA receptor neurotoxins through plant-insect coevolution. Nat Ecol Evol 7, 1444–1456 (2023). 10.1038/s41559-023-02127-4

9 Djoko Tagne, C. S. et al. A single mutation G454A in the P450 CYP9K1 drives pyrethroid resistance in the major malaria vector Anopheles funestus reducing bed net efficacy. Genetics 229, 1–40 (2025). 10.1093/genetics/iyae181

10 Ingham, V. A. et al. Integration of whole genome sequencing and transcriptomics reveals a complex picture of the reestablishment of insecticide resistance in the major malaria vector Anopheles coluzzii. PLoS Genet 17, e1009970 (2021). 10.1371/journal.pgen.1009970

11 Bass, C. et al. The evolution of insecticide resistance in the peach potato aphid, Myzus persicae. Insect Biochem Mol Biol 51, 41–51 (2014). 10.1016/j.ibmb.2014.05.003

12 Daborn, P. J. et al. A single p450 allele associated with insecticide resistance in Drosophila. Science 297, 2253–2256 (2002). 10.1126/science.1074170

13 Li, X., Schuler, M. A. & Berenbaum, M. R. Molecular mechanisms of metabolic resistance to synthetic and natural xenobiotics. Annu Rev Entomol 52, 231–253 (2007). 10.1146/annurev.ento.51.110104.151104

14 Homem, R. A. et al. Evolutionary trade-offs of insecticide resistance - The fitness costs associated with target-site mutations in the nAChR of Drosophila melanogaster. Mol Ecol 29, 2661–2675 (2020). 10.1111/mec.15503

15 Ffrench-Constant, R. H., Rocheleau, T. A., Steichen, J. C. & Chalmers, A. E. A point mutation in a Drosophila GABA receptor confers insecticide resistance. Nature 363, 449–451 (1993). 10.1038/363449a0

16 Mutero, A., Pralavorio, M., Bride, J. M. & Fournier, D. Resistance-associated point mutations in insecticide-insensitive acetylcholinesterase. Proc Natl Acad Sci U S A 91, 5922–5926 (1994). 10.1073/pnas.91.13.5922

17 McKenzie, J. A. & Batterham, P. Predicting insecticide resistance: mutagenesis, selection and response. Philos Trans R Soc Lond B Biol Sci 353, 1729–1734 (1998). 10.1098/rstb.1998.0325

18 Angst, D. C., Tepekule, B., Sun, L., Bogos, B. & Bonhoeffer, S. Comparing treatment strategies to reduce antibiotic resistance in an in vitro epidemiological setting. Proc Natl Acad Sci U S A 118 (2021). 10.1073/pnas.2023467118

19 Gatenby, R. A. & Brown, J. S. Integrating evolutionary dynamics into cancer therapy. Nat Rev Clin Oncol 17, 675–686 (2020). 10.1038/s41571-020-0411-1

20 Tabashnik, B. E. Managing resistance with multiple pesticide tactics: theory, evidence, and recommendations. J Econ Entomol 82, 1263–1269 (1989). 10.1093/jee/82.5.1263

21 REX Consortium. Heterogeneity of selection and the evolution of resistance. Trends Ecol Evol 28, 110–118 (2013). 10.1016/j.tree.2012.09.001

22 Cloyd, R. A. Pesticide mixtures and rotations: Are these viable resistance mitigating strategies. Pest Technology, 14–18 (2010).

23 Madgwick, P. G. & Kanitz, R. Beyond redundant kill: A fundamental explanation of how insecticide mixtures work for resistance management. Pest Manag Sci 79, 495–506 (2022). 10.1002/ps.7180

24 Denholm, I. & Rowland, M. W. Tactics for managing pesticide resistance in arthropods: theory and practice. Annu Rev Entomol 37, 91–112 (1992). 10.1146/annurev.en.37.010192.000515

25 Mani, G. S. Evolution of resistance in the presence of two insecticides. Genetics 109, 761–783 (1985). 10.1093/genetics/109.4.761

26 Caprio, M. A. Evaluating Resistance Management Strategies for Multiple Toxins in the Presence of External Refuges. Journal of Economic Entomology 91, 1021–1031 (1998). 10.1093/jee/91.5.1021

27 Roush, R. T. & McKenzie, J. A. Ecological genetics of insecticide and acaricide resistance. Annu Rev Entomol 32, 361–380 (1987). 10.1146/annurev.en.32.010187.002045

28 Roush, R. T. Designing resistance management programs: How can you choose? Pesticide Science 26, 423–441 (1989). 10.1002/ps.2780260409

29 Curtis, C. F. & Otoo, L. N. A simple model of the build-up of resistance to mixtures of anti-malarial drugs. Trans R Soc Trop Med Hyg 80, 889–892 (1986). 10.1016/0035-9203(86)90248-8

30 Comins, H. N. Tactics for resistance management using multiple pesticides. Agriculture, Ecosystems & Environment 16, 129–148 (1986). 10.1016/0167-8809(86)90099-X

31 Roush, R. T. Two-toxin strategies for management of insecticidal transgenic crops: can pyramiding succeed where pesticide mixtures have not? Philos Trans R Soc Lond B Biol Sci 353, 1777–1786 (1998). 10.1098/rstb.1998.0330

32 Curtis, C. F. Theoretical models of the use of insecticide mixtures for the management of resistance. Bulletin of Entomological Research 75, 259–266 (1985). 10.1017/S0007485300014346

33 Curtis, C. F. H. N. ; Kasim, S. H. Are there effective resistance management strategies for vectors of human disease? Biological Journal of the Linnean Society 48, 3–18 (1993). 10.1006/bijl.1993.1002

34 Tabashnik, B. E., Brevault, T. & Carriere, Y. Insect resistance to Bt crops: lessons from the first billion acres. Nat Biotechnol 31, 510–521 (2013). 10.1038/nbt.2597

35 Castle, S. J., Toscano, N. C., Prabhaker, N., Henneberry, T. J. & Palumbo, J. C. Field evaluation of different insecticide use strategies as resistance management and control tactics for Bemisia tabaci (Hemiptera: Aleyrodidae). Bull Entomol Res 92, 449–460 (2002). 10.1079/ber2002194

36 Immaraju, J. A. M., J. G. Hobza, R. F. Field Evaluation of Insecticide Rotation and Mixtures as Strategies for Citrus Thrips (Thysanoptera: Thripidae) Resistance Management in California. Journal of Economic Entomology 83, 306–314 (1990). 10.1093/jee/83.2.306

37 Zhao, J. Z. et al. Transgenic plants expressing two Bacillus thuringiensis toxins delay insect resistance evolution. Nat Biotechnol 21, 1493–1497 (2003). 10.1038/nbt907

38 Zhao, J. Z., Collins, H. L. & Shelton, A. M. Testing insecticide resistance management strategies: mosaic versus rotations. Pest Manag Sci 66, 1101–1105 (2010). 10.1002/ps.1985

39 Pimentel, D. & Burgess, M. Effects of Single Versus Combinations of Insecticides on the Development of Resistance. Environmental Entomology 14, 582–589 (1985). 10.1093/ee/14.5.582

40 Zoh, M. G. et al. Experimental evolution supports the potential of neonicotinoid-pyrethroid combination for managing insecticide resistance in malaria vectors. Sci Rep 11, 19501 (2021). 10.1038/s41598-021-99061-x

41 Sadia, C. G. et al. The impact of agrochemical pollutant mixtures on the selection of insecticide resistance in the malaria vector Anopheles gambiae: insights from experimental evolution and transcriptomics. Malar J 23, 69 (2024). 10.1186/s12936-023-04791-0

42 Macdonald, R. S., Surgeoner, G. A., Solomon, K. R. & Harris, C. R. Effect of Four Spray Regimes on the Development of Permethrin and Dichlorvos Resistance, in the Laboratory, by the House Fly (Diptera: Muscidae). Journal of Economic Entomology 76, 417–422 (1983). 10.1093/jee/76.3.417

43 McKenzie, C. L. & Byford, R. L. Continuous, alternating, and mixed insecticides affect development of resistance in the horn fly (Diptera: Muscidae). J Econ Entomol 86, 1040–1048 (1993). 10.1093/jee/86.4.1040

44 McGaughey, W. H. & Johnson, D. E. Indianmeal Moth (Lepidoptera: Pyralidae) Resistance to Different Strains and Mixtures of Bacillus thuringiensis. Journal of Economic Entomology 85, 1594–1600 (1992). 10.1093/jee/85.5.1594

45 Georghiou, G. P. & Wirth, M. C. Influence of Exposure to Single versus Multiple Toxins of Bacillus thuringiensis subsp. israelensis on Development of Resistance in the Mosquito Culex quinquefasciatus (Diptera: Culicidae). Appl Environ Microbiol 63, 1095–1101 (1997). 10.1128/aem.63.3.1095-1101.1997

46 Tabashnik, B. E. & McGaughey, W. H. Resistance Risk Assessment for Single and Multiple Insecticides: Responses of Indianmeal Moth (Lepidoptera: Pyralidae) to Bacillus thuringiensis. Journal of Economic Entomology 87, 834–841 (1994). 10.1093/jee/87.4.834

47 Burden, G. S., Lofgren, C. S. & Smith, C. N. Development of Chlordane and Malathion Resistance in the German Cockroach. Journal of Economic Entomology 53, 1138–1139 (1960). 10.1093/jee/53.6.1138

48 Fardisi, M., Gondhalekar, A. D., Ashbrook, A. R. & Scharf, M. E. Rapid evolutionary responses to insecticide resistance management interventions by the German cockroach (Blattella germanica L.). Sci Rep 9, 8292 (2019). 10.1038/s41598-019-44296-y

49 Parker, W. E., Howard, J. J., Foster, S. P. & Denholm, I. The effect of insecticide application sequences on the control and insecticide resistance status of the peach-potato aphid, Myzus persicae (Hemiptera:Aphididae), pon field crops of potato. Pest Manag Sci 62, 307–315 (2006). 10.1002/ps.1162

50 Prabhaker, N. T. N.,,,, Henneberry T. Evaluation of Insecticide Rotations and Mixtures as Resistance Management Strategies for Bemisia argentifolii (Homoptera: Aleyrodidae). Journal of Economic Entomology 91, 820–826 (1998). 10.1093/jee/91.4.820

51 Pimentel, D. & Bellotti, A. C. Parasite-host population systems and genetic stability. Am Nat 110 (1976). 10.1086/283110

52 Ffrench-Constant, R. H. The molecular genetics of insecticide resistance. Genetics 194, 807–815 (2013). 10.1534/genetics.112.141895

53 Grigoraki, L. et al. CRISPR/Cas9 modified An. gambiae carrying kdr mutation L1014F functionally validate its contribution in insecticide resistance and combined effect with metabolic enzymes. PLoS Genet 17, e1009556 (2021). 10.1371/journal.pgen.1009556

54 Essandoh, J., Yawson, A. E. & Weetman, D. Acetylcholinesterase (Ace-1) target site mutation 119S is strongly diagnostic of carbamate and organophosphate resistance in Anopheles gambiae s.s. and Anopheles coluzzii across southern Ghana. Malar J 12, 404 (2013). 10.1186/1475-2875-12-404

55 Wang, L. et al. Cadherin repeat 5 mutation associated with Bt resistance in a field-derived strain of pink bollworm. Sci Rep 10, 16840 (2020). 10.1038/s41598-020-74102-z

56 Jouraku, A. et al. Ryanodine receptor mutations (G4946E and I4790K) differentially responsible for diamide insecticide resistance in diamondback moth, Plutella xylostella L. Insect Biochem Mol Biol 118, 103308 (2020). 10.1016/j.ibmb.2019.103308

57 Schedl, T. & Kimble, J. fog-2, a germ-line-specific sex determination gene required for hermaphrodite spermatogenesis in Caenorhabditis elegans. Genetics 119, 43–61 (1988). 10.1093/genetics/119.1.43

58 Hu, S. et al. Multi-modal regulation of C. elegans hermaphrodite spermatogenesis by the GLD-1-FOG-2 complex. Dev Biol 446, 193–205 (2019). 10.1016/j.ydbio.2018.11.024

59 Guest, M., Kriek, N. & Flemming, A. J. Studies of an insecticidal inhibitor of acetyl-CoA carboxylase in the nematode C. elegans. Pestic Biochem Physiol 169, 104604 (2020). 10.1016/j.pestbp.2020.104604

60 Hahnel, S. R. et al. Extreme allelic heterogeneity at a Caenorhabditis elegans beta-tubulin locus explains natural resistance to benzimidazoles. PLoS Pathog 14, e1007226 (2018). 10.1371/journal.ppat.1007226

61 Roush, R. T. Occurrence, genetics and management of insecticide resistance. Parasitol Today 9, 174–179 (1993). 10.1016/0169-4758(93)90141-2

62 Kont, M. D. et al. Characterising the intensity of insecticide resistance: A novel framework for analysis of intensity bioassay data. Current Research in Parasitology & Vector-borne Diseases 4 (2023).

63 Tekoh, T. A. et al. The duplicated cytochrome P450 CYP6P9a/b confers cross-resistance to a mitochondrial complex I inhibitor in the African malaria vector Anopheles funestus. BMC Genomics 26, 837 (2025). 10.1186/s12864-025-11984-1

64 David, J. P., Ismail, H. M., Chandor-Proust, A. & Paine, M. J. Role of cytochrome P450s in insecticide resistance: impact on the control of mosquito-borne diseases and use of insecticides on Earth. Philos Trans R Soc Lond B Biol Sci 368, 20120429 (2013). 10.1098/rstb.2012.0429

65 Balabanidou, V., Grigoraki, L. & Vontas, J. Insect cuticle: a critical determinant of insecticide resistance. Curr Opin Insect Sci 27, 68–74 (2018). 10.1016/j.cois.2018.03.001

66 Li, L. Q. et al. A proof-of-concept experimental-theoretical model to predict pesticide resistance evolution. Heredity (Edinb) (2025). 10.1038/s41437-025-00781-x

67 Driscoll, M., Dean, E., Reilly, E., Bergholz, E. & Chalfie, M. Genetic and molecular analysis of a Caenorhabditis elegans beta-tubulin that conveys benzimidazole sensitivity. J Cell Biol 109, 2993–3003 (1989). 10.1083/jcb.109.6.2993

68 Gutbrod, P. et al. Inhibition of acetyl-CoA carboxylase by spirotetramat causes growth arrest and lipid depletion in nematodes. Sci Rep 10, 12710 (2020). 10.1038/s41598-020-69624-5

69 Fritz, M. L. et al. Contemporary evolution of a Lepidopteran species, Heliothis virescens, in response to modern agricultural practices. Mol Ecol 27, 167–181 (2018). 10.1111/mec.14430

70 West, S. A., Dall, S. R. X., Cunningham, J. P., Alonzo, S. H. & Griffin, A. S. Behavioural ecology in the twenty-first century. Nat Ecol Evol 9, 2193–2205 (2025). 10.1038/s41559-025-02912-3

71 Travis, J. & Reznick, D. N. Natural Selection: How Selection on Behavior Interacts with Selection on Morphology. Curr Biol 28, R882–R884 (2018). 10.1016/j.cub.2018.07.006

72 Kimura, M. On the probability of fixation of mutant genes in a population. Genetics 47, 713–719 (1962). 10.1093/genetics/47.6.713

73 Helps, J. C., Paveley, N. D., White, S. & van den Bosch, F. Determinants of optimal insecticide resistance management strategies. J Theor Biol 503, 110383 (2020). 10.1016/j.jtbi.2020.110383

74 Helps, J. C., Paveley, N. D. & van den Bosch, F. Identifying circumstances under which high insecticide dose increases or decreases resistance selection. J Theor Biol 428, 153–167 (2017). 10.1016/j.jtbi.2017.06.007

75 Levick, B., South, A. & Hastings, I. M. A Two-Locus Model of the Evolution of Insecticide Resistance to Inform and Optimise Public Health Insecticide Deployment Strategies. PLoS Comput Biol 13, e1005327 (2017). 10.1371/journal.pcbi.1005327

76 Sudo, M., Takahashi, D., Andow, D. A., Suzuki, Y. & Yamanaka, T. Optimal management strategy of insecticide resistance under various insect life histories: Heterogeneous timing of selection and interpatch dispersal. Evol Appl 11, 271–283 (2017). 10.1111/eva.12550

77 Barbosa, S., Kay, K., Chitnis, N. & Hastings, I. M. Modelling the impact of insecticide-based control interventions on the evolution of insecticide resistance and disease transmission. Parasit Vectors 11, 482 (2018). 10.1186/s13071-018-3025-z

78 South, A. & Hastings, I. M. Insecticide resistance evolution with mixtures and sequences: a model-based explanation. Malar J 17, 80 (2018). 10.1186/s12936-018-2203-y

79 Madgwick, P. G., Tunstall, T. & Kanitz, R. Evolutionary rescue in resistance to pesticides. Proc Biol Sci 291, 20240805 (2024). 10.1098/rspb.2024.0805

80 Madgwick, P. G. & Kanitz, R. Modelling new insecticide-treated bed nets for malaria-vector control: how to strategically manage resistance? Malar J 21, 102 (2022). 10.1186/s12936-022-04083-z

81 Hobbs, N. P., Weetman, D. & Hastings, I. M. Insecticide resistance management strategies for public health control of mosquitoes exhibiting polygenic resistance: A comparison of sequences, rotations, and mixtures. Evol Appl 16, 936–959 (2023). 10.1111/eva.13546

82 Martinello, M., Naggie, S., Rockstroh, J. K. & Matthews, G. V. Direct-Acting Antiviral Therapy for Treatment of Acute and Recent Hepatitis C Virus Infection: A Narrative Review. Clin Infect Dis 77, S238–S244 (2023). 10.1093/cid/ciad344

83 Nuwagaba, J., Li, J. A., Ngo, B. & Sutton, R. E. 30 years of HIV therapy: Current and future antiviral drug targets. Virology 603, 110362 (2025). 10.1016/j.virol.2024.110362

84 West, J. et al. Towards Multidrug Adaptive Therapy. Cancer Res 80, 1578–1589 (2020). 10.1158/0008-5472.CAN-19-2669

85 Labrie, M., Brugge, J. S., Mills, G. B. & Zervantonakis, I. K. Therapy resistance: opportunities created by adaptive responses to targeted therapies in cancer. Nat Rev Cancer 22, 323–339 (2022). 10.1038/s41568-022-00454-5

86 Laskey, S. B. & Siliciano, R. F. A mechanistic theory to explain the efficacy of antiretroviral therapy. Nat Rev Microbiol 12, 772–780 (2014). 10.1038/nrmicro3351

87 Perelson, A. S. & Deeks, S. G. Drug effectiveness explained: the mathematics of antiviral agents for HIV. Sci Transl Med 3, 91ps30 (2011). 10.1126/scitranslmed.3002656

88 Nyhoegen, C., Bonhoeffer, S. & Uecker, H. The many dimensions of combination therapy: How to combine antibiotics to limit resistance evolution. Evol Appl 17, e13764 (2024). 10.1111/eva.13764

89 Pomeroy, A. E., Schmidt, E. V., Sorger, P. K. & Palmer, A. C. Drug independence and the curability of cancer by combination chemotherapy. Trends Cancer 8, 915–929 (2022). 10.1016/j.trecan.2022.06.009

90 Zhang, J., Cunningham, J., Brown, J. & Gatenby, R. Evolution-based mathematical models significantly prolong response to abiraterone in metastatic castrate-resistant prostate cancer and identify strategies to further improve outcomes. Elife 11 (2022). 10.7554/eLife.76284

91 Otto, S. P. & Lenormand, T. Resolving the paradox of sex and recombination. Nat Rev Genet 3, 252–261 (2002). 10.1038/nrg761

92 Becks, L. & Agrawal, A. F. The evolution of sex is favoured during adaptation to new environments. PLoS Biol 10, e1001317 (2012). 10.1371/journal.pbio.1001317

93 Roze, D. & Barton, N. H. The Hill-Robertson effect and the evolution of recombination. Genetics 173, 1793–1811 (2006). 10.1534/genetics.106.058586

94 Hartley, C. J. et al. Amplification of DNA from preserved specimens shows blowflies were preadapted for the rapid evolution of insecticide resistance. Proc Natl Acad Sci U S A 103, 8757–8762 (2006). 10.1073/pnas.0509590103

95 Toprak, E. et al. Evolutionary paths to antibiotic resistance under dynamically sustained drug selection. Nat Genet 44, 101–105 (2011). 10.1038/ng.1034

96 Smith, J. M. & Haigh, J. The hitch-hiking effect of a favourable gene. Genet Res 23, 23–35 (1974).

97 Greaves, M. & Maley, C. C. Clonal evolution in cancer. Nature 481, 306–313 (2012). 10.1038/nature10762

98 Desai, M. M. & Fisher, D. S. Beneficial mutation selection balance and the effect of linkage on positive selection. Genetics 176, 1759–1798 (2007). 10.1534/genetics.106.067678

99 Sulston, J. & Hodgkin, J. in The Nematode Caenorhabditis elegans (ed B Wood) (Cold Spring Harbor Laboratory Press, 1988).

100 Crow, J. F. & Kimura, M. An Introduction to Population Genetics Theory. (Harper & Row, 1970).

