## Supplementary material for "Constraining pesticide resistance using evolution-informed selection regimes": All supplementary figures and legends

**Extended Data Figures**

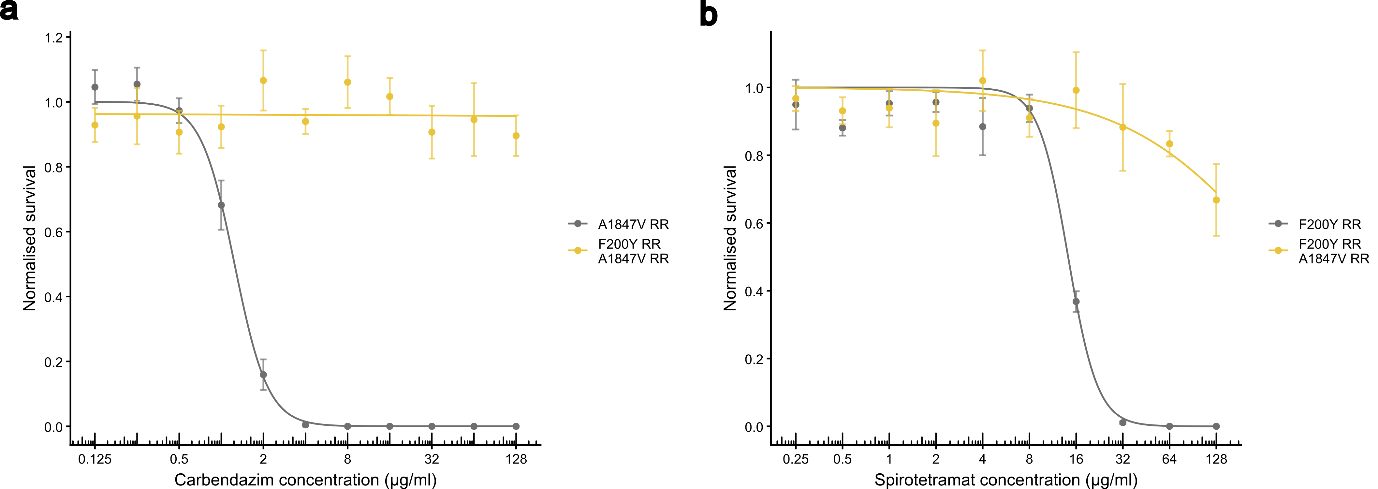

**Extended Data Figure 1. Dose-response analysis of cross-resistance and compound interaction effects.** a) Dose-survival curves showing the effect of harbouring homozygous *pod-2* A1847V alleles on carbendazim resistance on otherwise WT and carbendazim-resistant backgrounds. b) Dose-survival curves showing the effect of harbouring homozygous *ben-1* F200Y alleles on spirotetramat resistance on otherwise WT and spirotetramat-resistant backgrounds. Eggs were seeded onto compound wells on day 0, and survival was quantified as the number of healthy adults after 96h incubation at 20°C. Data represent control-normalised mean ± SEM from four independent biological replicates (*n* = 4). Curves were fitted using nonlinear regression (two-parameter logistic model) where possible, otherwise a linear model was fitted when no appreciable fitness decline could be observed at the highest compound concentrations.

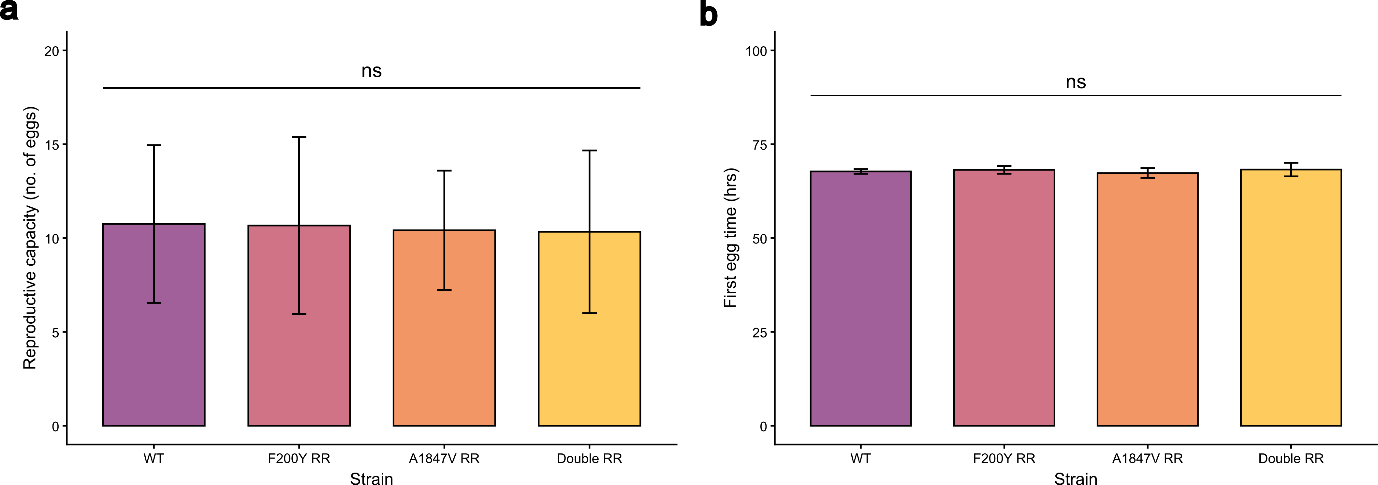

**Extended Data Figure 2. Early-life fitness analysis of WT and homozygous resistant strains.** a) Reproductive capacity, as quantified by the number of eggs carried by a mature adult female. Data represent mean ± SD from 12 adult females (n = 12). All resistant strains were compared to the WT and none showed a significant difference in reproductive capacity. b) Developmental rate, as quantified by number of hours between seeding F0 eggs and the appearance of the first F1 egg. Data represent mean ± SD from 12 biological replicates (*n* = 12). All resistant strains were compared to the WT and none showed a significant difference in developmental rate.

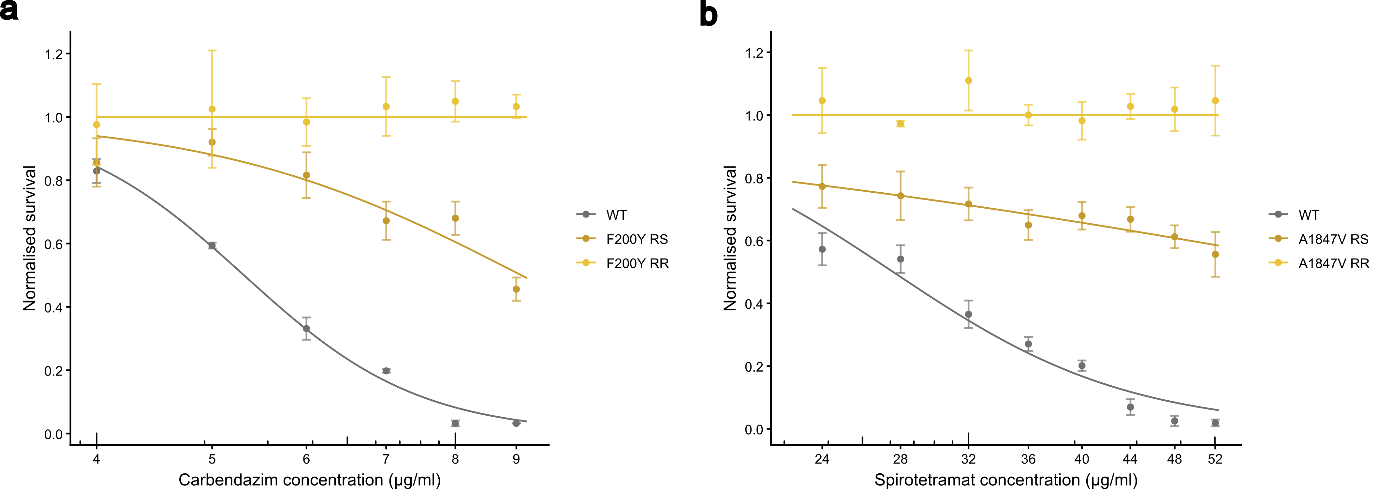

**Extended Data Figure 3. Dose-response analysis of resistance alleles upon 2-day compound exposure.** a) Dose-survival curves showing the effect of harbouring 0, 1, 2 copies of the *ben-1* F200Y allele on resistance to carbendazim. b) Dose-survival curves showing the effect of harbouring 0, 1, 2 copies of the *pod-2* A1847V allele on resistance to spirotetramat. Eggs were seeded onto compound wells on day 0, and transferred to no-compound plates on day 2. Survival was quantified as the number of healthy adults after a total of 96h indication at 20°C. Data represent control-normalised mean ± SEM from three independent biological replicates (*n* = 3). Curves for WT and heterozygotes were fitted using nonlinear regression (two-parameter logistic model). Homozygote-resistant individuals showed no evidence of decline in fitness at all tested compound concentrations.

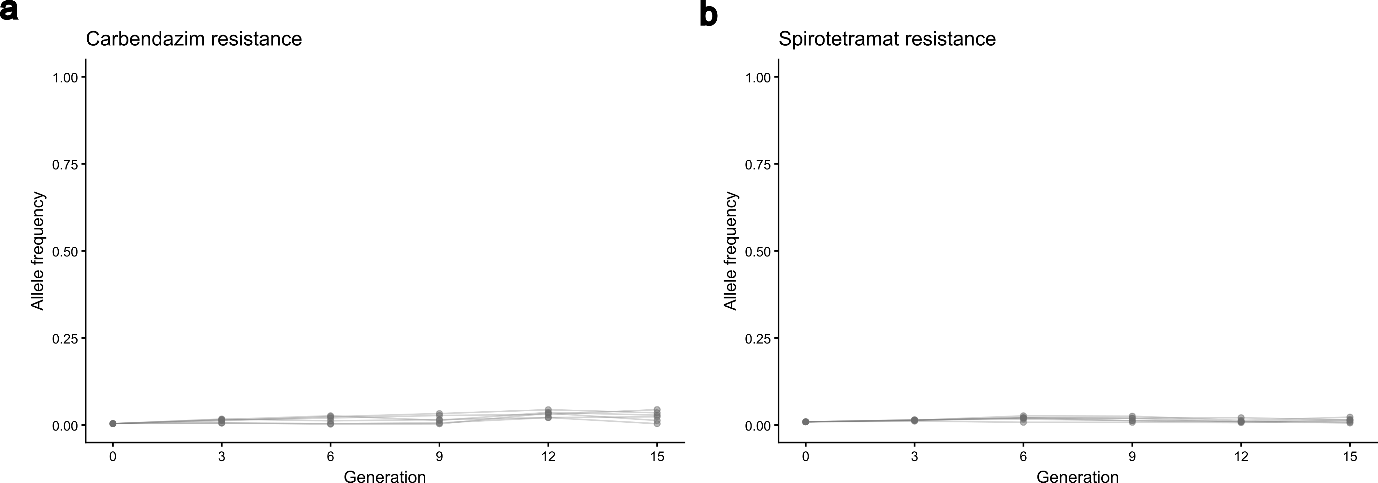

**Extended Data Figure 4. Resistance allele frequency dynamics under no selection.** a) Change in frequency of the carbendazim resistance allele. b) Change in frequency of the spirotetramat resistance allele. Allele frequencies were measured at the end of every three generations using high-depth Illumina amplicon sequencing. Data represent individual biological replicate trajectories across 15 generations of experimental evolution.

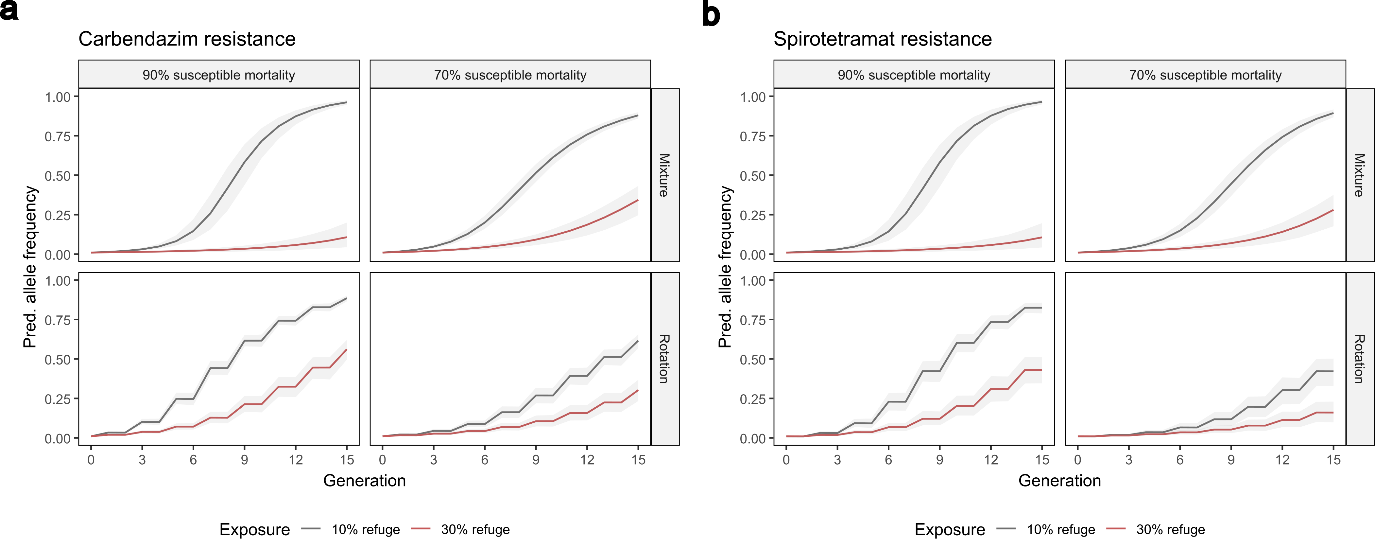

**Extended Data Figure 5. Computationally predicted resistance allele frequency dynamics under different selection regimes.** a) Change in frequency of the carbendazim resistance allele. b) Change in frequency of the spirotetramat resistance allele. 500 stochastic simulations were run for each distinct parameter combination. Data represent mean (coloured line) and the 2.5^th^ and 97.5^th^ percentile range (ribbon).
