## Supplementary table 1 for "Constraining pesticide resistance using evolution-informed selection regimes"

| Genetic locus | Primer name | Sequence (5’ → 3’) |
| --- | --- | --- |
| *ben-1* | ben-1 F1 | AGTCATTGAGCGCTACTCTTTCTGTCCACCA |
| *ben-1* | ben-1 F2 | GATCTCATTCCGCTACTCTTTCTGTCCACCA |
| *ben-1* | ben-1 F3 | CGCTTATCCTCGCTACTCTTTCTGTCCACCA |
| *ben-1* | ben-1 F4 | CGACGTAGTCCGCTACTCTTTCTGTCCACCA |
| *ben-1* | ben-1 F5 | TCAATGATCGCGCTACTCTTTCTGTCCACCA |
| *ben-1* | ben-1 F6 | AGTCTCGGCACGCTACTCTTTCTGTCCACCA |
| *ben-1* | ben-1 R1 | ATCTGCGTACTGGTGACTCCGGACATTGTAAC |
| *ben-1* | ben-1 R2 | GATTGCACGCTGGTGACTCCGGACATTGTAAC |
| *ben-1* | ben-1 R3 | ATGCTTCCTATGGTGACTCCGGACATTGTAAC |
| *ben-1* | ben-1 R4 | TGCTAACTTCTGGTGACTCCGGACATTGTAAC |
| *ben-1* | ben-1 R5 | ATAGCAGTGCTGGTGACTCCGGACATTGTAAC |
| *pod-2* | pod-2 F1 | GAGCGAGTCAAGTTCGGAGCACAGATCGTC |
| *pod-2* | pod-2 F2 | CAGGCGATCTAGTTCGGAGCACAGATCGTC |
| *pod-2* | pod-2 F3 | TTCACGGAAGAGTTCGGAGCACAGATCGTC |
| *pod-2* | pod-2 F4 | GAGCAGATATAGTTCGGAGCACAGATCGTC |
| *pod-2* | pod-2 F5 | CGGCTACAGTAGTTCGGAGCACAGATCGTC |
| *pod-2* | pod-2 F6 | ACGATGAAGTAGTTCGGAGCACAGATCGTC |
| *pod-2* | pod-2 R1 | GTAAGGCTCCCTCCATCATCATTGGCTTGCG |
| *pod-2* | pod-2 R2 | CCTCGAATGGCTCCATCATCATTGGCTTGCG |
| *pod-2* | pod-2 R3 | TCATGGAATCCTCCATCATCATTGGCTTGCG |
| *pod-2* | pod-2 R4 | ATAAGAGGTCCTCCATCATCATTGGCTTGCG |
| *pod-2* | pod-2 R5 | GCTGACCTGACTCCATCATCATTGGCTTGCG |

Table S1. Barcoding scheme for multiplexed amplicon sequencing. Six distinct forward and five distinct reverse PCR amplification primers were constructed for each genetic locus, resulting in a total of up to 30 unique combinations. Each sequencing primer begins with a unique 10bp barcode, followed by 20-22bp of a common sequence to selectively amplify ~220bp around the genetic locus of interest.
